# Age-related differences in the role of the prefrontal cortex in sensory-motor training gains: A tDCS study

**DOI:** 10.1101/2020.09.09.282285

**Authors:** Si Jing Tan, Hannah L. Filmer, Paul E. Dux

## Abstract

The ability to process multiple sources of information concurrently is particularly impaired as individuals age and such age-related increases in multitasking costs have been linked to impairments in response selection. Previous neuroimaging studies with young adults have implicated the left hemisphere prefrontal cortex (PFC) as a key neural substrate of response selection. In addition, several transcranial direct current stimulation studies (tDCS) have provided causal evidence implicating this region in response selection and multitasking operations. For example, Filmer at al. (2013b) demonstrated that typically observed response selection learning/training gains in young adults were disrupted via offline transcranial direct current stimulation (tDCS) of left, but not right, PFC. Here, considering evidence of functional dedifferentiation in the brains of older adults, we assessed if this pattern of response selection learning disruption via tDCS to the left PFC is observed in older adults, testing if this region remains a key response selection node as individuals age. In a pre-registered study with 58 older adults, we applied anodal, cathodal, and sham stimulation to left and right PFC, and measured performance as participants trained on low- and high-response selection load tasks. Active stimulation did not disrupt training in older adults as compared to younger adults. However, there was evidence of enhanced training gains via tDCS, which scaled with response selection task difficulty. The results highlight age-related differences in the casual neural substrates that subserve response selection and learning.

## Introduction

Ageing is associated with changes across multiple cognitive domains including processing speed, attention, memory, aspects of language, visuospatial abilities, and executive functioning (Harada et al., 2013). Of importance, higher-level cognitive processes such as executive functions show disproportionate age-related decline relative to other psychological operations (Kievit et al., 2014; Persson et al., 2016). Executive functions (EFs; also called executive control or cognitive control) refer to a collection of cognitive operations that are involved in selecting, planning, maintaining, and implementing goal-directed behaviour (Diamond, 2013) and are thought to be critical predictors of functional abilities and independence in older adults (Grigsby et al., 1998). Indeed, poorer executive functioning in older adults has been associated with increased risk of falls (Faulkner et al., 2007; Kearney et al., 2013; Mirelman et al., 2012) and lower driving competence (Daigneault et al., 2002).

Every-day functional activities require an individual to make goal-directed decisions where sensory information from the environment is mapped to the appropriate motor output – *response selection*. Given the overwhelming variety and amount of information humans are presented with, performance costs are inevitable. This is especially the case when individuals are time-pressured, have many decision options and/or are required to make two or more decisions simultaneously (i.e. multitasking (Garner and Dux, 2015)). Interestingly, such response selection and multitasking costs are observed even when tasks require relatively simple mappings despite the immense processing power of the brain (Marois and Ivanoff, 2005; Pashler, 1994).

There is extensive evidence indicating that older adults have more difficulty in carrying out two tasks concurrently relative to younger adults (Hartley and Little, 1999; Verhaeghen, 2011; Verhaeghen et al., 2003; Verhaeghen and Cerella, 2002). In simple choice reaction tasks, Sanders (1998) described six underlying stages of information processing, namely: pre-processing, feature extraction, identification, response selection, motor programming, and motor adjustment (Sanders, 2013). It has been suggested that the greatest deterioration in multitasking performance with age is predominantly attributable to changes in basic response selection processes - where stimulus properties are mapped to a motor output (Allen et al., 1998; Meiran et al., 2001; Meiran and Gotler, 2001; Melis et al., 2002; Salthouse and Miles, 2002; Stelmach et al., 1987). Furthermore, higher level cognitive operations are found to become more strongly reliant on lower-level sensory and motor mechanisms as individuals age (Li et al., 2001; Li and Dinse, 2002).

Response selection is often studied using multitask paradigms, such as in the psychological refractory period (PRP) (Dux et al., 2009, 2006; Lien et al., 2006; Pashler, 1994, 1984; Sigman and Dehaene, 2008; Welford, 1952), or under single-task conditions where response alternative load is manipulated (Filmer et al., 2013a). In a typical dual-task/PRP paradigm, two simple sensory-motor tasks are presented in quick succession. Typically, participants respond more slowly to the second of two tasks as the time interval between them is reduced. Several prominent explanations have been proposed to account for this task 2 slowing. The first being a structural “central bottleneck” framework where response selection processes are hypothesised to be capacity limited and only conducted serially (Pashler, 1994; Welford, 1952) causing task 2 to have to wait until task 1 response selection is complete. Conversely, Meyer and Kieras (1997a; 1997b) hypothesised that two response selection operations can be conducted in parallel once sensory-motor production rules are converted from declarative knowledge to procedural knowledge. In this theory, task 2 slowing occurs due to task structure and the system aiming to limit inter-task interference. Finally, central capacity sharing models (Navon and Miller, 2002; Tombu and Jolicoæur, 2003) posit that while response selection processes are capacity limited, there is some sharing of resources between the two tasks at the response selection stage. Of key relevance to the present work, the PRP is more pronounced in older adults, suggesting that there may be age-related changes specific to response selection. For example, even after accounting for generalised slowing, older adults have been found to be less efficient coordinating response selection processes across tasks (Allen et al., 1998) or, at the very least, may be using different task coordination strategies as compared to younger adults (Glass et al., 2000).

Neuroimaging studies have provided evidence that the posterior lateral prefrontal cortex (pLPFC) of the left hemisphere is a key neural substrate for this central bottleneck (Dux et al., 2009, 2006; Hesselmann et al., 2011; Ivanoff et al., 2009; Jiang and Kanwisher, 2003; Sigman and Dehaene, 2008; Tombu et al., 2011). For example, pLPFC showed a temporal profile of blood-oxygen-level-dependent (BOLD) activity that mapped onto task 2 slowing in a both a PRP paradigm and response selection load task (Dux et al., 2006). In addition, training on dual tasks was associated with reduced differences between single and dual task activation in the left pLPFC (Dux et al., 2009) mirroring reductions in RT.

More recently, transcranial direct current stimulation (tDCS) studies have demonstrated a causal relationship between the PFC and response selection (Filmer et al., 2013b, 2013a). tDCS is a non-invasive brain stimulation method that involves passing a weak electric current between two electrodes placed on the scalp. It is thought to influence resting membrane potentials and thus the likelihood of neural firing and so can be used to causally modulate cortical activity by applying either excitatory (e.g., anodal) or inhibitory (e.g., cathodal) stimulation protocols (Filmer et al., 2014). However, the neurophysiological basis of stimulation effects is likely more complex than a simple excitation/inhibition model (Filmer et al., 2020). In terms of response selection, Filmer and colleagues (2013b) showed that performance gains in response selection training/learning (i.e. decreased reaction times) were disrupted via offline stimulation (both anodal and cathodal) relative to sham in the left but not right PFC (Filmer et al., 2013b). This showed that the left PFC plays a key role in learning sensory-motor mappings, and this finding has since been replicated in a pre-registered study (Filmer et al., 2019). Studies have also shown improved multitasking performance following stimulation to the left PFC following a single session of tDCS (Filmer et al., 2013a) as well as stimulation combined with response-selection training across multiple sessions (Filmer et al., 2017).

The studies described above were all conducted with healthy younger adults; thus, it is yet to be determined if these results apply to older adults and if the same neural substrates are causally involved in response selection operations in this population. Indeed, healthy ageing has been associated with both increases (“overactivation”) and decreases (“underactivation”) in brain activation relative to younger adults, particularly in the prefrontal regions. “Overactivation” of regions has been associated with maintained performance and has been related to neural inefficiency, dedifferentiation of brain regions and compensatory processes (Carp et al., 2010; Park et al., 2004). Conversely, “underactivation” is often related to performance deficits, especially when more resources were required with increasing task complexity (Bennett et al., 2013; Cappell et al., 2010). In relation to multitasking, a previous study showed that older adults had increased brain activation in the left ventral PFC though decreased activation in left and right dorsolateral PFC following multitasking training (Erickson et al., 2007). Further to this, in a fMRI study directly comparing the neural substrates in dual-task interference between older and young adults, Hartley and colleagues (2011) suggested that both age groups showed similar underlying neural substrates involving a medial prefrontal and lateral frontal-parietal network. However, older adults showed additional recruitment of the frontopolar prefrontal cortex of which increased activation was observed to be correlated with successful reduction in dual-task interference (Hartley et al., 2011).

Collectively, there is now clear evidence of age-related changes in the neural substrates underlying higher cognitive processes. However, to date, there are few studies exploring the causal neural substrates that subserve response selection in older adults. Developing treatments for delaying the onset of executive function impairments associated with ageing (enhancing cognition) will rely on isolating key changes in brain regions/network across the lifespan that support high-level cognition. To that end, here we adopted the approach of Filmer et al. (2013b) to investigate the causal role of the left PFC in response selection and response selection learning/training in older adults. Specifically, the study examined if a similar pattern of results is observed (i.e. disruption of performance with active stimulation) for older adults, assessing if the left prefrontal cortex remains a key node for response selection as humans age (Filmer et al., 2013b).

## Material and methods

### Participants

A total of 58 participants were recruited. All but four were right-handed and reported normal (or corrected to normal) vision and hearing. All participants passed a tDCS safety-screening questionnaire and had no history of psychiatric, neurological or cognitive impairments. Four participants failed to meet the inclusion criteria (i.e. task performance > 60%) and were excluded from the final analysis. 27 (mean age = 69 years, range = 62 – 75 years, 16 females) participated in the left hemisphere condition, and 27 (mean age = 69 years, range = 62 – 75 years, 14 females) participated in the right hemisphere condition. Participants were stratified by demographic variables (i.e. age, education, gender), level of physical activity scores, and baseline executive functioning scores (see Table 1, for summary of measures). A script written in MatLab was used to allocate participants to hemisphere based on aforementioned factors, while concurrently accounting for the size of each group to ensure that allocation over time was approximately equal. Both conditions were conducted concurrently. The University of Queensland Human Research Ethics Committee approved the study and all participants provided informed, written consent. Participants were reimbursed $20 per hour of participation. The method and analyses (unless otherwise stated) were pre-registered (https://osf.io/8ksjd/).

**Table 1:**
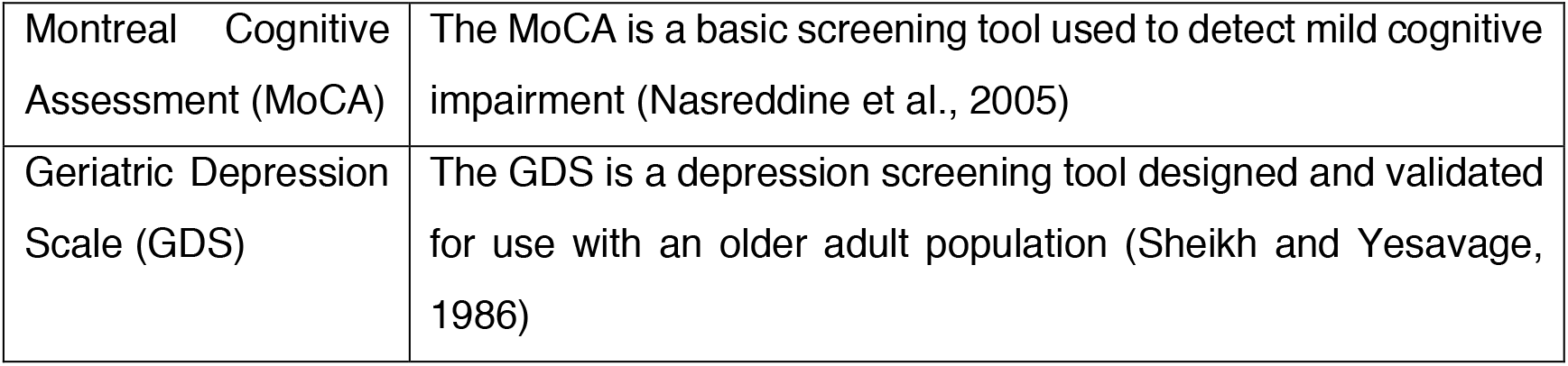
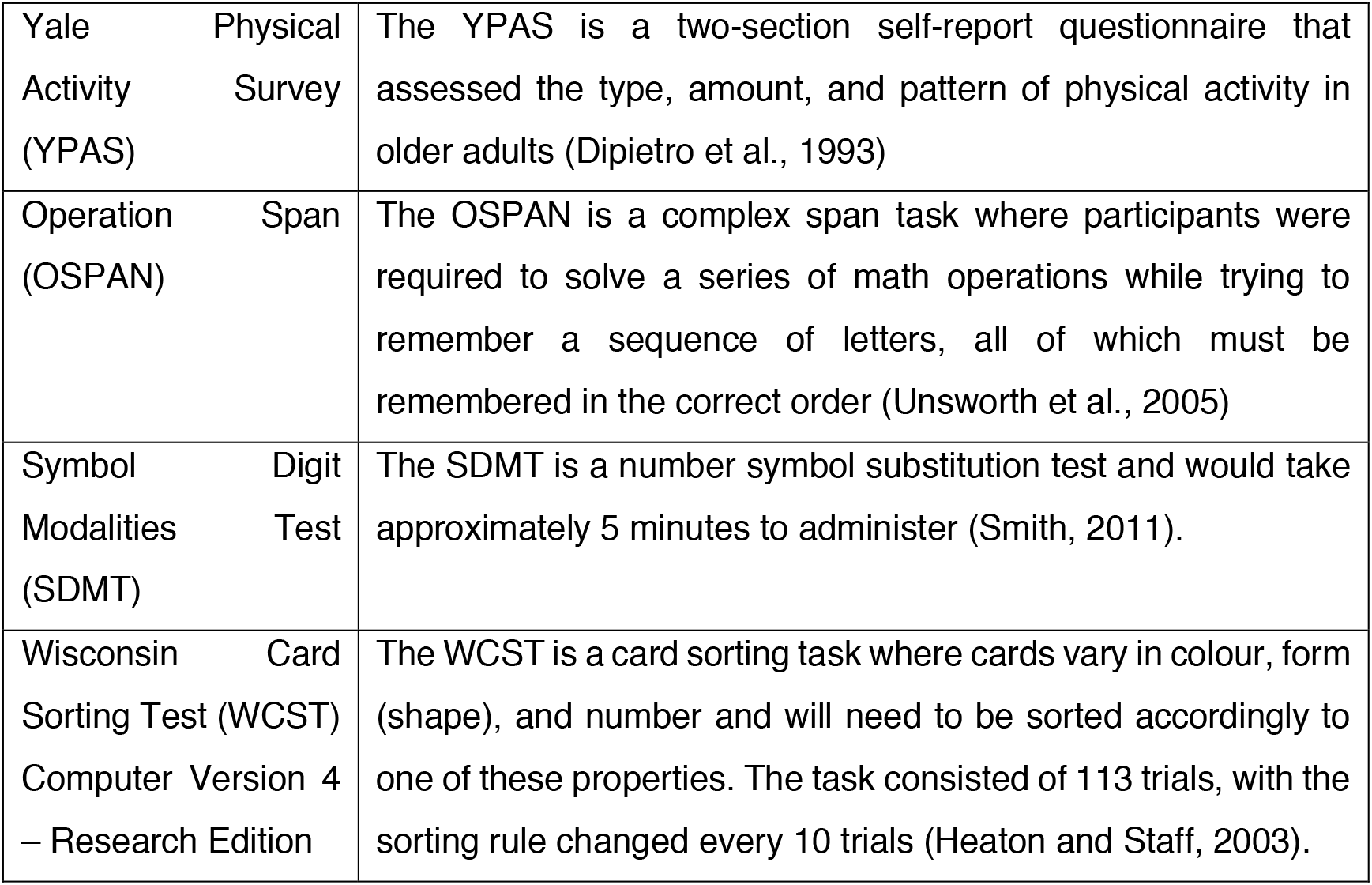
Summary of measures.

### Overview

Participants attended a total of four sessions. In the first session, they completed screening tests, questionnaires and a range of cognitive tasks, which were used to assign individuals to groups (see Table 1). Participants that met all inclusion criteria then proceeded on to the following three sessions. In the second, third and fourth sessions, participants completed a computerised simple choice reaction time task where they had to discriminate between stimuli. There were three different types of stimuli (i.e. coloured circles, symbols or complex tones) employed for the task, with one set of stimuli assigned to each session. This was done to avoid any cross-session practice effects and counter-balanced across participants. There was a total of six stimuli for each task set - two of the six stimuli were used for the low response load and the remaining four were used for the high response load. The high load condition consisted of fewer response alternatives (4 rather than 6) than were employed in younger adults (Filmer et al., 2013b) as it was anticipated, based on preliminary work in our lab, that the task would be more difficult for older adults to learn. As we were focussed on the effect of PFC stimulation, we attempted to equate baseline response selection difficulty between the groups. Participants completed 18 blocks of 30 trials that were separated into three phases of 6 blocks each (i.e. pre-tDCS, post-tDCS and 20mins post-tDCS). tDCS stimulation was applied after the pre-tDCS phase, with a different type of stimulation used in each session (anodal, cathodal, sham). To wit, the within-subjects variables for the experiment were task type (stimulus set), response load, training phase, and stimulation type, which are detailed further below. The between-subjects variable was brain region: left hemisphere vs. right hemisphere PFC.

### Stimulation protocol

The stimulation protocol replicated that of Filmer et al. (2013b). Each participant completed three stimulation sessions, with a different type of stimulation used each time (anodal, cathodal, sham). The pairing of stimulation type to session was controlled across participants, so each type of stimulation occurred equally often in each session (using a latin square). The sessions were conducted a minimum of 48 hours apart.

A battery-operated NeuroConn stimulator with two 5 × 5 cm electrodes was used to administer the current. The placement of each electrodes on the scalp was based on the 10 – 20 EEG system (Jasper, 1958). The positioning of the target electrode was 1cm posterior to the F3 electrode side for the left hemisphere and similarly 1cm posterior to the F4 electrode site for the right hemisphere. The reference electrode was placed over the contralateral supraorbital region ((Filmer et al., 2013b); see Figure 1). The stimulation intensity was 0.7mA for a total of 9 minutes for both the anodal and cathodal stimulation sessions (including 30 second ramp-up/ramp-down periods). The sham condition involved the same parameters but only for stimulation duration of 1 minute and 15 seconds (including 30 second ramp-up/ramp-down periods). The participants were told to sit quietly with their eyes open during the 9-minute stimulation and were blinded to the type of stimulation that they received.

**Figure 1:**
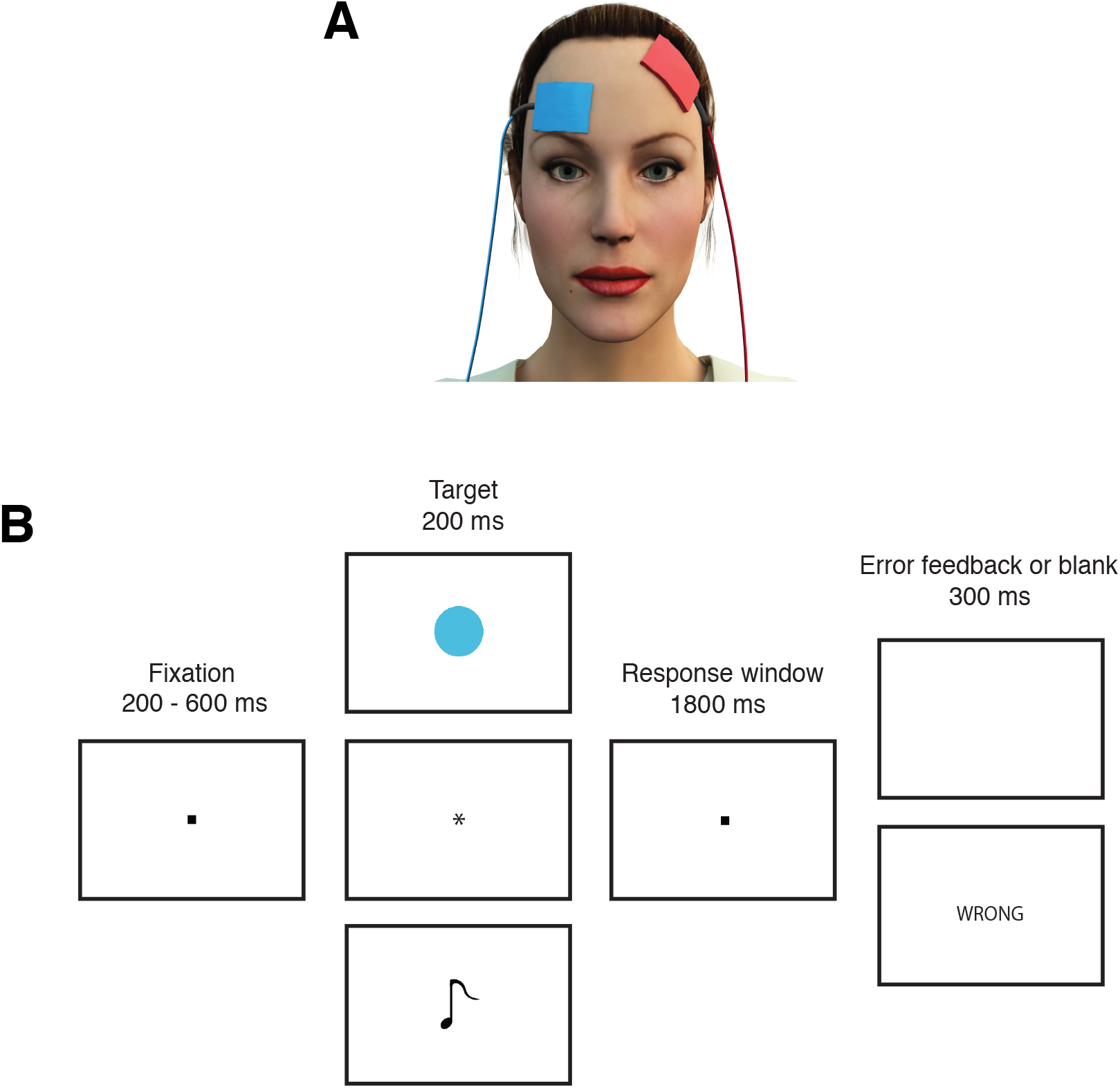
A) tDCS electrode montage. B) trial outline. Participants were shown a fixation dot on a monitor, which was followed by a stimulus (i.e. coloured circle, symbol, sound). They were instructed to respond as quickly and as accurately as possible. Participants received either an error feedback or a blank screen (accurate response) depending on their responses.

### Behavioural task

Participants were asked to distinguish between sets of either coloured circles (dark green: RGB 10 130 65, light green: RGB 109 205 119, light blue: RGB 79 188 220, brown: RGB 167 106 48, pink: RGB 255 57 255, and yellow: RGB 255 235 30), computer symbols (%, #, ~, @, *, *) or complex tones (see (Filmer et al., 2013b)). Each experimental session used one task, and the these was counterbalanced across the participants and paired to experimental sessions equally often. Each task was used only once in order for separate training/learning effects to be observed in all three stimulation sessions. Each task consisted of a low response load condition (two possible stimuli-responses) and high response load condition (four possible stimuli-responses). Half of the participants responded with the index fingers of both hands for the low response load, and the ring and middle of both hands for the high response load. The other half of the participants responded with the ring finger of both hands for the low response load, and the middle and index fingers of both hands of the high response load. This was to counterbalance any difference in dexterity of fingers. Participants had a repeating block pattern of high response load followed up with low response load, and vice versa for the other half of participants. Three practice blocks of 30 trials for each response selection load condition were given to participants for familiarisation and accuracy feedback (see Figure 1). During the experiment, participants were seated 70cm from a 24” ASUS monitor with a refresh rate of 144Hz. Participants were shown a fixation dot located centrally on the monitor for 200ms – 600ms. This was followed by a target stimulus (i.e. a coloured circle, a symbol or a complex tone) presented for 200ms. Complex tones were played through Bose headphones that participants wore throughout the experiment. Volume was adjusted when necessary. Participants were instructed to respond as quickly and as accurately as possible. No response after 1800ms, or an inaccurate response, resulted in error feedback. A minimum of 60% accuracy was required to move on to the main experiment.

Following familiarisation, participants completed eighteen blocks of 30 trials, which was divided into three phases (six blocks per phase). tDCS was applied (offline) after the first phase. Reaction times were measured at three time points: before tDCS stimulation, immediately after tDCS stimulation (i.e. second phase completed immediately after the 9 minutes stimulation) and 20 minutes after the cessation of stimulation.

### Analysis

For this study, we employed both Bayesian and Frequentist statistical approaches implemented in JASP (JASP Team, 2019) to analyse the data. Interpretations were based on the Bayesian findings, with frequentist included given their prominence in the literature. All analyses were pre-registered (https://osf.io/8ksjd/) unless described as exploratory.

We used a standard interpretation of Bayes factors (van Doorn et al., 2019), where a Bayes factor of 1 indicate the same amount of evidence for both the null (H_0_) and alternative hypothesis (H_1_). Bayes factors between 1 and 3 were considered to be weak or inconclusive evidence for the alternative hypothesis. Bayes factors between 3 and 10 were considered moderate evidence while Bayes factors greater than 10 were considered strong evidence for the alternative hypothesis (H_1_). On the contrary, Bayes factors between 0.33 and 1 were considered to be weak or inconclusive evidence for the null hypothesis (H_0_). Bayes factors between 0.10 and 0.33 were considered moderate evidence while Bayes factors less than 0.03 were considered strong evidence for the null hypothesis (H_0_).

Large single-task learning/training improvements occur rapidly in such RT paradigms as used here (Dux et al., 2006), thus the key question of interest concerned how training/learning effects, within session (i.e. difference between pre-tDCS and 20 minutes post-tDCS phase), as measured with accuracy and reaction times (RTs), were impacted by stimulation polarity, stimulation hemisphere, and response selection load.

### Data availability

All study materials and data are publicly available.

## Results

### Reaction Times (RTs)

#### Impact of training on RTs

Participants showed training-related performance gains (i.e. reduction in RTs) across time for both response loads and in all three stimulation sessions. RTs for each response load, in each hemisphere are shown in Figures 2-3.

**Figure 2:**
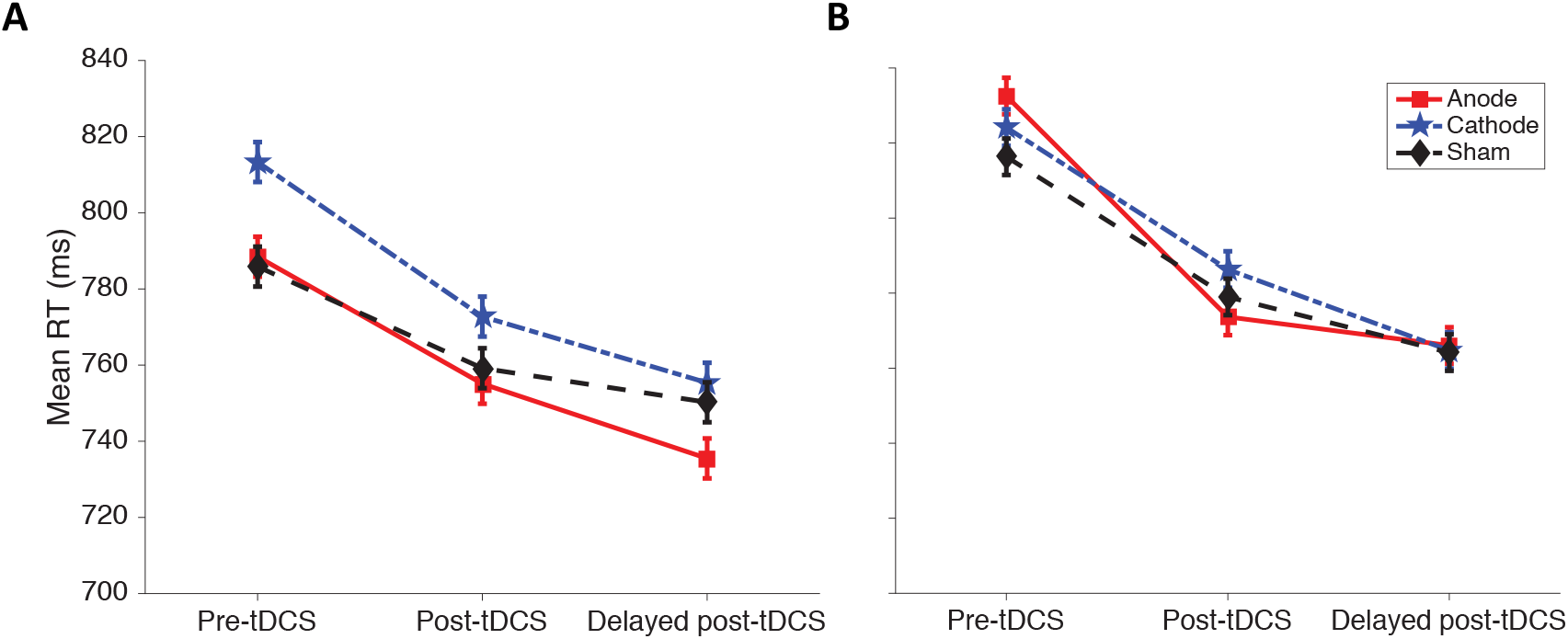
Impact of training and stimulation on high response load trials. A) and B) show the average RT for each stimulation session across the three phases of the experiment for the left and right hemisphere conditions, respectively. Error bars for all graphs represent SEM of the change in RT compared with the pre-tDCS phase.

**Figure 3.**
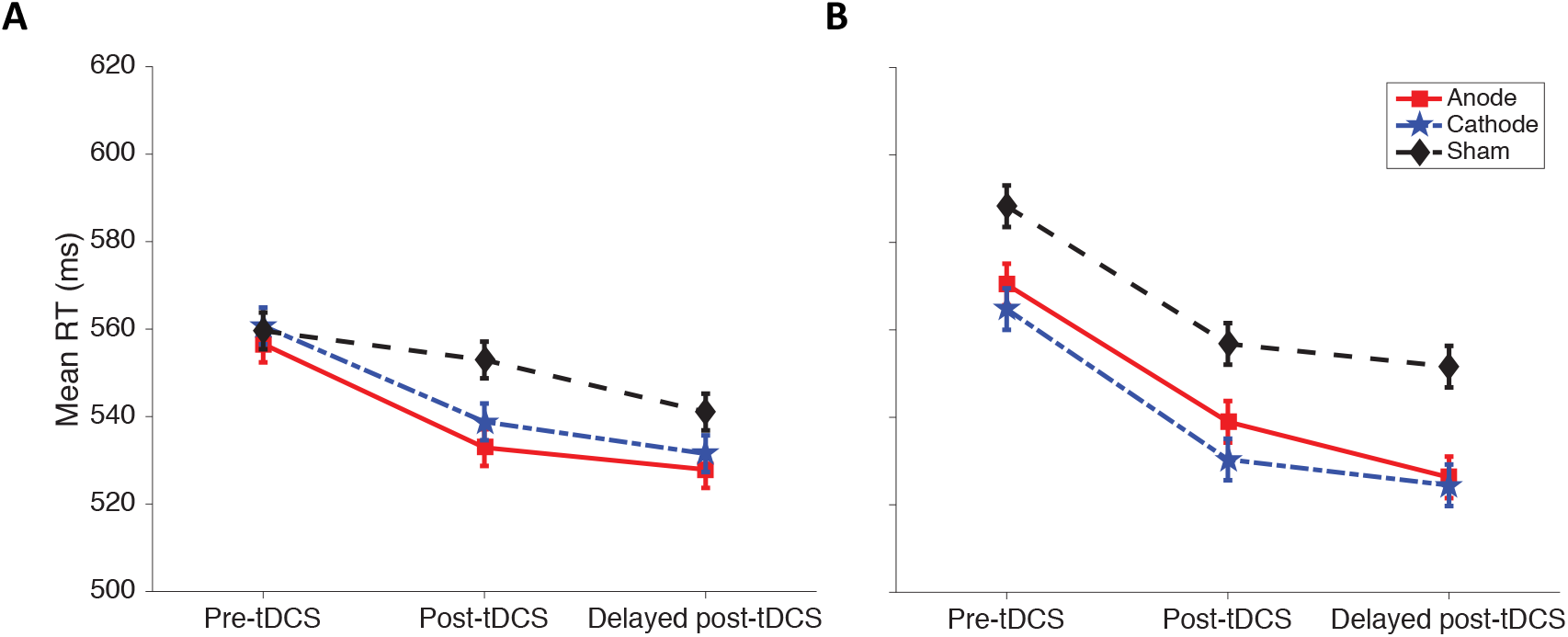
Impact of training and stimulation on low response load. A) and B) show the average RT for each stimulation session across the three phases of the experiment for the left and right hemisphere conditions, respectively. Error bars for all graphs represent SEM of the change in RT compared with the pre-tDCS phase.

In the left hemisphere, for high response load condition, there was strong evidence for training-related performance gains for the anode (BF_10_ = 1701.40; *t*(26) = 5.37, *p* < 0.001), cathode (BF_10_ = 7504.35; *t*(26) = 5.99, *p* < 0.001) and sham sessions (BF_10_ = 617.78; *t*(26) = 4.94, *p* < 0.001). For the low response selection load, there was strong evidence for training-related performance gains for the anode (BF_10_ = 25.41; *t*(26) = 3.58, *p* = 0.001) and cathode sessions (BF_10_ = 227.51; *t*(26) = 4.52, *p* < 0.001) and weak evidence for the sham session (BF_10_ = 2.91; *t*(26) = 2.54, *p* = 0.018).

Similarly in the right-hemisphere, participants showed training-related performance gains (i.e. reduction in RTs) across time in both response loads. For the high response load, there was strong evidence for anode (BF_10_ = 43574.69; *t*(26) = 6.74, *p* < 0.001), cathode (BF_10_ = 178657.30; *t*(26) = 7.35, *p* < 0.001), and sham sessions (BF_10_ = 147218.36; *t*(26) = 7.27, *p* < 0.001). Similarly, for the low response load, strong evidence for training-related performance gains was found for anode (BF_10_ = 466.05; *t*(26) = 4.82, *p* < 0.001), cathode (BF_10_ = 3765.79; *t*(26) = 5.70, *p* < 0.001), and sham sessions (BF_10_ = 127.78; *t*(26) = 4.28, *p* < 0.001).

The greater reaction time training effects for the high compared to low response load, seen across both hemisphere, suggest that our response load manipulation tapped the central bottleneck. Specifically, for the left hemisphere, there were strong evidence (BF_10_ = 65.88; *t*(26) = 3.99, *p* < 0.001) when comparing training effects in the high (*M* = 0.05, *SD* = 0.03) to the low response selection load (*M* = 0.03, *SD* = 0.03;). For the right hemisphere, there were moderate evidence (BF_10_ = 5.21; *t*(26) = 2.83, *p* = 0.009) when comparing training effects in the high (*M* = 0.06, *SD* = 0.03) to the low (*M* = 0.04, *SD* = 0.03) response load. Notably, low response load training benefits were only observed in our older adult study not observed in our previous young adult study (Filmer et al., 2013b).

#### Impact of stimulation of RTs

Planned paired samples t-tests were used to examine differences in training effects in the high response load (i.e. difference in RTs between pre-tDCS and 20mins post-tDCS) between the three stimulation sessions, separately for the left and the right hemisphere conditions.

Overall, for the left hemisphere, there was no evidence of meaningful differences in training effects between the active and sham sessions for the high response load. There was weak evidence for the null hypothesis when comparing training improvements between anode and sham sessions (BF_10_ = 0.86; *t*(26) = 1.82, *p* = 0.080), and cathode and sham sessions (BF_10_ = 0.90; *t*(26) = 1.85, *p* = 0.075).

Similarly, there were no meaningful differences in training effects in active versus sham stimulation for the high response load in the right hemisphere. There was weak evidence for the null hypothesis when comparing between anode and sham sessions (BF_10_ = 0.37; *t*(26) = 1.16, *p* = 0.256), and moderate evidence for the null hypothesis when comparing cathode and sham sessions (BF_10_ = 0.26; *t*(26) = 0.74, *p* = 0.468).

### Response Accuracy

Participants had high response accuracy across all the phases, load and hemisphere conditions. Accuracy data can be found in Table 2. There were no effects of stimulation on response accuracy across both hemispheres (BF_10_ < 1 – 0.1). Overall, there appeared to be no speed/accuracy trade-offs, although it should be noted accuracy was close to ceiling.

**Table 2.** Mean accuracy rates for all conditions.

| **ACC (%)** | **Left** |  |  | **Right** |  |  |
| --- | --- | --- | --- | --- | --- | --- |
|  | **Before tDCS** | **Immediate post** | **20mins post** | **Before tDCS** | **Immediate post** | **20mins post** |
| **Anode** |  |  |  |  |  |  |
| High RS | 95.49 | 97.42 | 97.99 | 95.95 | 98.03 | 98.30 |
| Low RS | 98.34 | 98.73 | 98.53 | 97.42 | 98.69 | 98.61 |
| **Cathode** |  |  |  |  |  |  |
| High RS | 93.87 | 96.95 | 96.91 | 95.37 | 96.53 | 97.61 |
| Low RS | 97.53 | 97.57 | 98.42 | 98.80 | 98.80 | 98.92 |
| **Sham** |  |  |  |  |  |  |
| High RS | 97.11 | 97.88 | 97.80 | 96.60 | 97.45 | 97.15 |
| Low RS | 98.11 | 97.99 | 98.65 | 98.34 | 98.65 | 98.57 |

### Exploratory analyses

#### Training effects of task types and stimulations conditions

Response selection is thought to be an central process and thus not dependent stimulus modality (Dux et al., 2009, 2006; Ivanoff et al., 2009; Jiang and Kanwisher, 2003; Pashler, 1994, 1984; Welford, 1952). However, in the present study we did observed that different sets of stimuli were associated with different levels of difficulty. In particular, within the high response load condition, baseline pre-stimulation RTs in the sound discrimination task were higher (*M* = 921ms) than those in the circle discrimination task (*M* = 750ms) and symbol discrimination task (*M* = 760ms).

To examine the presence of any possible difficulty and modality effects, we separated the RT results into the three different task types for the high response load trials. Analyses were then conducted to compare the training effects across task types and stimulation conditions. The results suggest a facilitation effect of active stimulation relative to sham that varied across task type.

##### Sounds discrimination task

Overall, the results show that there are greater training effects for the sound discrimination task in both hemispheres, following active relative to sham stimulation (see Figure 4). Specifically, in the left hemisphere, comparing anode and sham sessions, there was strong evidence for enhanced training outcomes (faster responses in the 20 mins post vs pre phase; BF_10_ = 26.98; *t*(8) = 4.64, *p* = 0.002). A comparison of cathode and sham sessions revealed similarly strong evidence for enhanced training outcomes (BF_10_ = 11.92; *t*(8) = 3.90, *p* = 0.005; see Figure 4).

**Figure 4.**
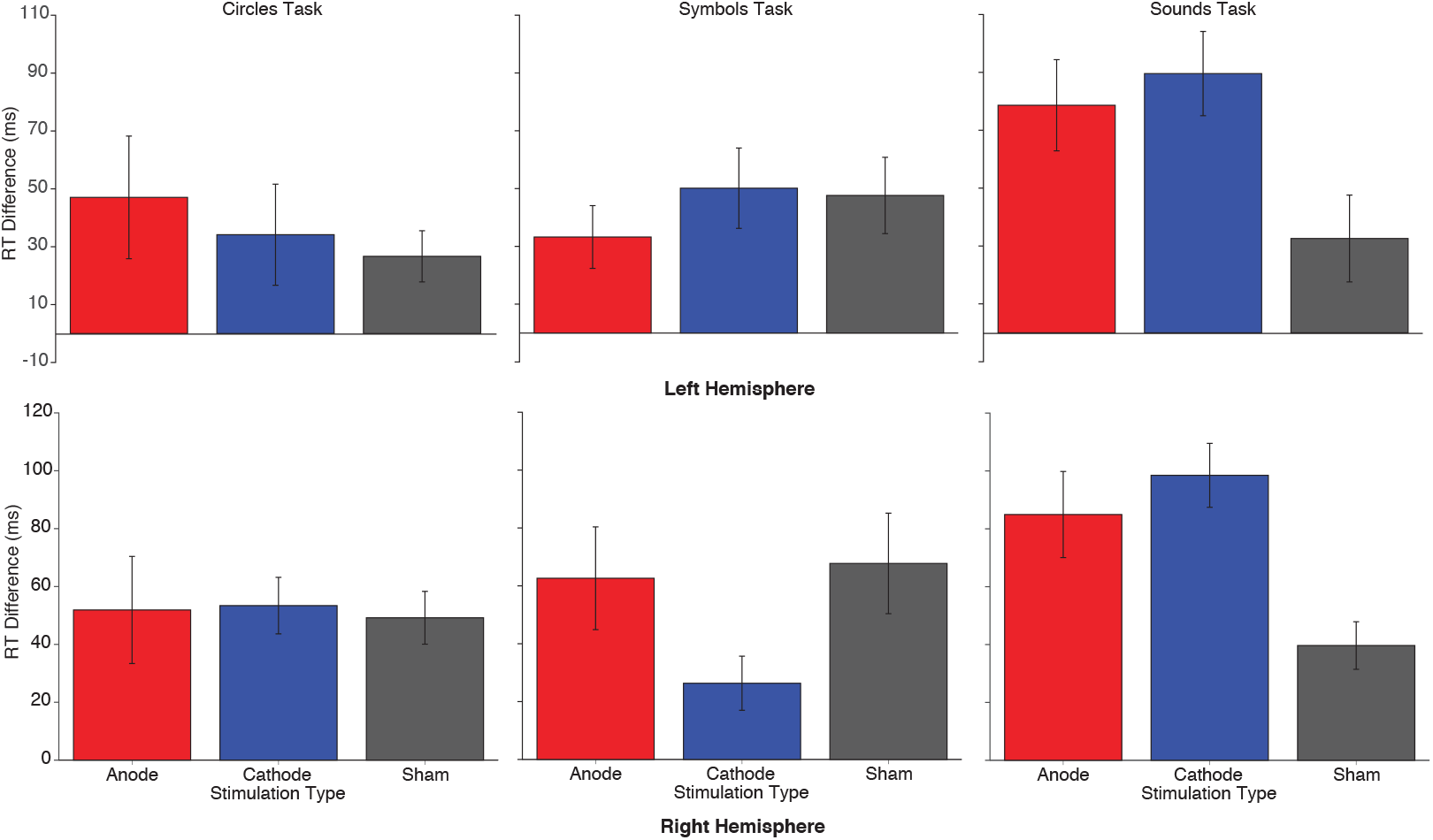
Effect of left hemisphere (above) and right hemisphere (below) stimulation on training for the high response load task across three stimulus types.

In the right hemisphere, comparing anode and sham sessions, there was moderate evidence for enhanced training outcomes (BF_10_ = 3.68; *t*(8) = 2.90, *p* = 0.020). A comparison of cathode and sham sessions showed strong evidence for enhanced training outcomes (BF_10_ = 15.06; *t*(8) = 4.11, *p* = 0.003; see Figure 4).

##### Symbols discrimination task

Overall, there was little evidence for stimulation modulating training outcomes for the symbols task. For the left hemisphere, all effects were weakly in favour of the null (BF_10_ between 0.33 – 1; see Figure 4).

For the right hemisphere, comparison between anode and sham stimulation was also weakly in favour of the null (BF_10_ between 0.33 - 1). However, there was weak evidence for enhanced training outcomes when comparing cathode to anode sessions (BF_10_ = 1.22; *t*(8) = −1.95, *p* = 0.086) as well as cathode to sham sessions (BF_10_ = 1.15; *t*(8) = −1.90, *p* = 0.094; see Figure 4).

##### Colours discrimination task

Across both hemispheres, all tests comparing active relative to sham sessions did not show any difference in training effects (BF_10_ between 0.33 – 1; see Figure 4).

#### Comparison of older and young adults

In Filmer et al. (2013b), active stimulation (anodal and cathodal) disrupted performance gains for younger adults in the high response load condition following stimulation to the left PFC but not to the right PFC (Filmer et al., 2013b). The results suggested that the left PFC plays a causal role in response selection (i.e. learning sensory-motor mappings) for young adults. Here, we investigated the differences in the neural substrates of response selection between older and young adults, in the left hemisphere only, by comparing the present results to those of Filmer et al (2013b).

Analyses were first conducted to determine if there were baseline differences in pre-stimulation RTs between age groups across stimulation sessions. There was no difference in pre-stimulation RTs across the high load condition between age groups indicated by moderate evidence for the null when comparing cathode (BF_10_ = 0.31; *t*(43) = −0.24, *p* = 0.813) and anode sessions (BF_10_ = 0.31; *t*(43) = −1.47, *p* = 0.148). There was inconclusive evidence for differences in pre-stimulation RTs between age groups for sham session (BF_10_ = 1.75; *t*(43) = −2.12, *p* = 0.040). Indeed, older adults had slightly faster pre-stimulation RTs (*M* = 0.79, *SD* = 0.13) compared to young adults (*M* = 0.88, *SD* = 0.16) in the sham session. Note, however, that older adults in our study were presented with 4 alternative forced choices (AFC) in the high response load compared to the 6AFC in the young adult study. Thus, our approach for equating difficulty across the age-groups was successful.

There was a significant difference of stimulation on training effects between the age groups. The two-way interaction between training effects (i.e. difference in RTs between pre-tDCS and 20mins post-tDCS) and stimulation – training effects x stimulation – with between subjects factors of age showed strong evidence for the alternative hypothesis (BF_10_ = 75.93, *F*(1.64, 70.45) = 7.46, *p* < 0.002). This illustrated that stimulation had a different effect on performance gains when comparing across age groups. Specifically, active stimulation disrupted training-related performance (i.e. reaction times) gains relative to sham sessions in younger adults, but not in older adults. Follow-up independent samples t-tests comparing both age groups were conducted. There was strong evidence for age-related differences in training effects both for the cathode sessions (BF_10_ = 45.89; *t*(43) = 3.69, *p* < 0.001) and moderate evidence for anode sessions (BF_10_ = 6.64; *t*(43) = 2.85, *p* = 0.007, see Figure 5). In this context, our results suggest age-related changes in the neural substrates that underlie response selection and response selection learning.

**Figure 5:**
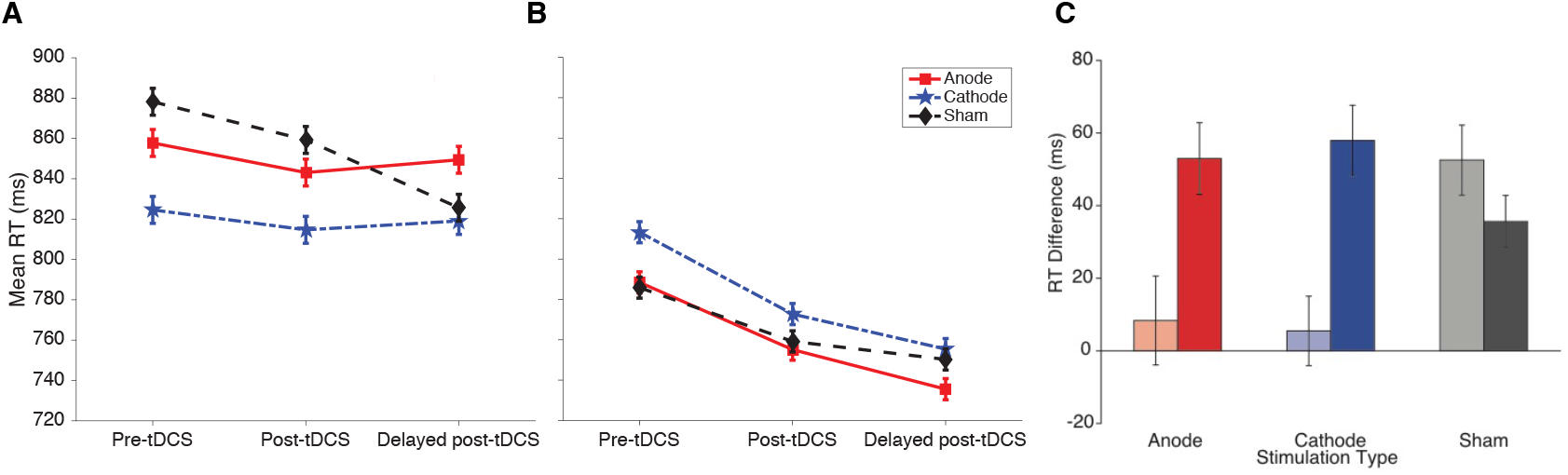
Impact of training and stimulation on high response load (left hemisphere) trials. A) and B) show the RT for each stimulation sessions for each of the three phases of the experiment for the younger A) and older adults B). C) shows the comparison of training effects of RTs in high response load between younger and older adults (left hemisphere). Lighter shades represent younger adults while darker shades represent older adults.

## Discussion

Here we investigated the causal role of the left PFC in response selection and response selection learning/training in older adults, testing if this region remains a key structure in this cognitive operation as individual age. We applied tDCS (active (anodal or cathodal) vs. sham stimulation) to both the left and right PFC midway through response selection training/learning. Participants trained on low- (2 alternate forced choice) and high- (4 alternate forced choice) response load tasks. Active stimulation of both the left and right hemisphere PFC did not disrupt response selection training performance. This is in contrast to previous data with young adults which showed that active stimulation of the left PFC disrupted performance gains (i.e. decreased RTs) under high response load conditions. Taking the young adult data into context, our findings are suggestive of age-related changes in the neural substrates of response selection with the left PFC no longer being a specific node for response selection with advancing age.

There was very little evidence for baseline RTs differences across age-groups. Training effects were present for the older adult group in the sham stimulation condition, indicating that they benefitted from response selection training as did the younger cohort. Older adults had greater performance gains with training (i.e. decreased RTs) for the high-versus low-response load, showing that our manipulation tapped the response selection bottleneck. This is the case even though there was a reduced number of forced choice in the high-load condition for the older adults (4AFC) compared to the young adults (6AFC, see (Filmer et al., 2013b)). Thus, this step to equate baseline performance across all conditions and equate training effect patterns between the ages appears to have been successful. It is also worthy to note that despite the load effects, older adults still found the low response load task difficult and displayed training effects, which is not the case for younger adults (Filmer et al., 2013b).

A general explanation for the present results, based on a speed/accuracy trade-off, was also considered. There is no evidence of a trade-off but to note, older adults were already performing at high accuracy rates in the beginning of the experiment, which could have masked an effect. In addition, and of import, we controlled for variability in education, age, gender, and baseline executive functioning scores and allocated participants approximately equally across both hemisphere conditions, thus it is unlikely these factors impacted on the results. Finally, our study recruitment criteria only included older adults that passed a cognitive screening test, and stimulation effects are reflective of age-related changes in healthy older adults and not attributable to an underlying neurodegenerative process.

Interestingly, exploratory analyses revealed stimulation had dissociable effects based on the type of task older adults completed. Both anodal and cathodal stimulation of the PFC, relative to sham for the high response load showed enhancement of performance for the most difficult task that subjects completed – the auditory sensory-motor training task – across *both* left and right hemisphere. This diffuse bilateral pattern of enhancement was not observed in our young adult sample. It is unlikely that it was due to age-related sensory changes as this facilitation effect was not observed in the low load condition. Thus, our results show preliminary evidence that in older adult’s response selection training gains can be improved via the modulation of PFC activity, which indicates that limitations of response selection processes in this group may be attenuated via such interventions.

Most studies in the literature utilising tDCS with cognitive training in older adults tend to use only anodal tDCS relative to sham over the dorsolateral prefrontal cortex, as cathodal stimulation has been generally linked to disruption of performance (Jones et al., 2015; Park et al., 2014; Stephens and Berryhill, 2016). Our polarity non-specific facilitation effects of tDCS specific to the auditory sensory-motor task show preliminary results that tDCS have far more complex impacts on the aging brain and would require further investigations.

Lastly, in context of cognitive aging, our results reflect a reduction in regional specialisation or specificity in particular areas, lending support to the idea of de-differentiation in an ageing brain. As facilitation effects were seen with both anodal and cathodal stimulation, it is difficult to ascertain if this was due compensatory mechanisms. Future studies on response selection need to consider using functional magnetic resonance imaging (fMRI) or electroencephalogram (EEG) concurrently with tDCS to investigate the effects of performance and task-related brain activity modulated by tDCS.

## Conclusions

Our findings suggest that the left PFC ceases to play a specific role in response selection learning/training with age. In comparison to a young adult population, two main differences were found. Active stimulation (i.e. anodal and cathodal) applied to the left PFC did not disrupt response selection training performance gains. In addition, performance enhancement via tDCS to both left and right PFC for auditory sensory-motor learning (the most difficult task subjects undertook) was observed. Our findings provide the first causal evidence of age-related changes in the neural substrates of response selection.

## Notes

**Funding**: HLF was supported by a UQ Fellowship (UQFEL1607881) and ARC Discovery Early Career Researcher Award (DE190100299).

https://osf.io/8ksjd/

### Competing Interest Statement

The authors have declared no competing interest.

## References

Allen, P.A., Smith, A.F., Vires-Collins, H., Sperry, S., 1998. The psychological refractory period: Evidence for age differences in attentional time-sharing. Psychol. Aging 13, 218–229. https://doi.org/10.1037/0882-7974.13.2.218

Bennett, I.J., Rivera, H.G., Rypma, B., 2013. Isolating age-group differences in working memory load-related neural activity: Assessing the contribution of working memory capacity using a partial-trial fMRI method. Neuroimage 72, 20–32. https://doi.org/10.1016/j.neuroimage.2013.01.030

Cappell, K.A., Gmeindl, L., Reuter-Lorenz, P.A., 2010. Age differences in prefontal recruitment during verbal working memory maintenance depend on memory load. Cortex 46, 462–473. https://doi.org/10.1016/j.cortex.2009.11.009

Carp, J., Gmeindl, L., Reuter-Lorenz, P.A., 2010. Age differences in the neural representation of working memory revealed by multi-voxel pattern analysis. Front. Hum. Neurosci. 4, 1–10. https://doi.org/10.3389/fnhum.2010.00217

Daigneault, G., Joly, P., Frigon, J.Y., 2002. Executive functions in the evaluation of accident risk of older drivers. J. Clin. Exp. Neuropsychol. 24, 221–238. https://doi.org/10.1076/jcen.24.2.221.993

Diamond, A., 2013. Executive Functions. Annu. Rev. Psychol. 64, 135–168. https://doi.org/10.1146/annurev-psych-113011-143750

Dipietro, L., Caspersen, C.J., Ostfeld, A.M., Nadel, E.R., 1993. A survey for assessing physical activity among older adults. Med. Sci. Sports Exerc. 25, 628–42.

Dux, P.E., Ivanoff, J., Asplund, C.L., Marois, R., 2006. Isolation of a Central Bottleneck of Information Processing with Time-Resolved fMRI. Neuron 52, 1109–1120. https://doi.org/10.1016/j.neuron.2006.11.009

Dux, P.E., Tombu, M.N., Harrison, S., Rogers, B.P., Tong, F., Marois, R., 2009. Training improves multitasking performance by increasing the speed of information processing in human prefrontal cortex. Neuron 63, 127–138. https://doi.org/10.1016/j.neuron.2009.06.005

Erickson, K.I., Colcombe, S.J., Wadhwa, R., Bherer, L., Peterson, M.S., Scalf, P.E., Kim, J.S., Alvarado, M., Kramer, A.F., 2007. Training-induced plasticity in older adults: Effects of training on hemispheric asymmetry. Neurobiol. Aging 28, 272–283. https://doi.org/10.1016/j.neurobiolaging.2005.12.012

Faulkner, K.A., Redfern, M.S., Cauley, J.A., Landsittel, D.P., Studenski, S.A., Rosano, C., Simonsick, E.M., Harris, T.B., Shorr, R.I., Ayonayon, H.N., Newman, A.B., 2007. Multitasking: Association Between Poorer Performance and a History of Recurrent Falls. J Am Geriatr Soc 55, 570–576. https://doi.org/10.1111/j.1532-5415.2007.01147.x

Filmer, H.L., Dux, P.E., Mattingley, J.B., 2014. Applications of transcranial direct current stimulation for understanding brain function. Trends Neurosci. 37, 742–753. https://doi.org/10.1016/j.tins.2014.08.003

Filmer, H.L., Ehrhardt, S.E., Bollmann, S., Mattingley, J.B., Dux, P.E., 2019. Accounting for individual differences in the response to tDCS with baseline levels of neurochemical excitability. Cortex 115, 324–334. https://doi.org/10.1016/j.cortex.2019.02.012

Filmer, H.L., Mattingley, J.B., Dux, P.E., 2020. Modulating brain activity and behaviour with tDCS: Rumours of its death have been greatly exaggerated. Cortex 123, 141–151. https://doi.org/10.1016/j.cortex.2019.10.006

Filmer, H.L., Mattingley, J.B., Dux, P.E., 2013a. Improved multitasking following prefrontal tDCS. Cortex 49, 2845–2852. https://doi.org/10.1016/j.cortex.2013.08.015

Filmer, H.L., Mattingley, J.B., Marois, R., Dux, P.E., 2013b. Disrupting prefrontal cortex prevents performance gains from sensory-motor training. J. Neurosci. 33, 18654–18660. https://doi.org/10.1523/JNEUROSCI.2019-13.2013

Filmer, H.L., Varghese, E., Hawkins, G.E., Mattingley, J.B., Dux, P.E., 2017. Improvements in Attention and Decision-Making Following Combined Behavioral Training and Brain Stimulation. Cereb. Cortex 27, 3675–3682. https://doi.org/10.1093/cercor/bhw189

Garner, K.G., Dux, P.E., 2015. Training conquers multitasking costs by dividing task representations in the frontoparietalsubcortical system. Proc. Natl. Acad. Sci. U. S. A. 112, 14372–14377. https://doi.org/10.1073/pnas.1511423112

Glass, J.M., Schumacher, E.H., Lauber, E.J., Zurbriggen, E.L., Gmeindl, L., Kieras, D.E., Meyer, D.E., 2000. Aging and the psychological refractory period: Task-coordination strategies in young and old adults. Psychol. Aging 15, 571–595. https://doi.org/10.1037/0882-7974.15.4.571

Grigsby, J., Kaye, K., Baxter, J., Shetterly, S.M., Hamman, R.F., 1998. Executive cognitive abilities and functional status among community-dwelling older persons in the San Luis Valley health and aging study. J. Am. Geriatr. Soc. 46, 590–596. https://doi.org/10.1111/j.1532-5415.1998.tb01075.x

Harada, C.N., Natelson Love, M.C., Triebel, K.L., 2013. Normal cognitive aging. Clin. Geriatr. Med. 29, 737–752. https://doi.org/10.1016/j.cger.2013.07.002

Hartley, A.A., Jonides, J., Sylvester, C.Y.C., 2011. Dual-task processing in younger and older adults: Similarities and differences revealed by fMRI. Brain Cogn. 75, 281–291. https://doi.org/10.1016/j.bandc.2011.01.004

Hartley, A.A., Little, D.M., 1999. Age-related differences and similarities in dual-task interference. J. Exp. Psychol. Gen. 128, 416–449. https://doi.org/10.1037/0096-3445.128.4.416

Heaton, R., Staff, P., 2003. Wisconsin Card Sorting Test®: Computer Version 4–Research Edition [WWW Document]. Psychol. Assess. Resour. URL https://www.parinc.com/Products/Pkey/483

Hesselmann, G., Flandin, G., Dehaene, S., 2011. Probing the cortical network underlying the psychological refractory period: A combined EEG-fMRI study. Neuroimage 56, 1608–1621. https://doi.org/10.1016/j.neuroimage.2011.03.017

Ivanoff, J., Branning, P., Marois, R., 2009. Mapping the pathways of information processing from sensation to action in four distinct sensorimotor tasks. Hum. Brain Mapp. 30, 4167–4186. https://doi.org/10.1002/hbm.20837

Jasper, H., 1958. The ten twenty electrode system of the international federation. Electroencephalogr. Clin. Neurophysiol. 10, 371–375.

Jiang, Y., Kanwisher, N., 2003. Common Neural Substrates for Response Selection across Modalities and Mapping Paradigms. J. Cogn. Neurosci. 15, 1080–1094. https://doi.org/10.1162/089892903322598067

Jones, K.T., Stephens, J.A., Alam, M., Bikson, M., Berryhill, M.E., 2015. Longitudinal neurostimulation in older adults improves working memory. PLoS One 10, 1–18. https://doi.org/10.1371/journal.pone.0121904

Kearney, F.C., Harwood, R.H., Gladman, J.R.F., Lincoln, N., Masud, T., 2013. The relationship between executive function and falls and gait abnormalities in older adults: A systematic review. Dement. Geriatr. Cogn. Disord. 36, 20–35. https://doi.org/10.1159/000350031

Kievit, R.A., Davis, S.W., Mitchell, D.J., Taylor, J.R., Duncan, J., Henson, R.N.A., 2014. Distinct aspects of frontal lobe structure mediate age-related differences in fluid intelligence and multitasking. Nat. Commun. 5, 1–10. https://doi.org/10.1038/ncomms6658

Li, S.C., Dinse, H.R., 2002. Aging of the brain, sensorimotor, and cognitive processes. Neurosci. Biobehav. Rev. https://doi.org/10.1016/S0149-7634(02)00059-3

Li, S.C., Lindenberger, U., Sikström, S., 2001. Aging cognition: From neuromodulation to representation. Trends Cogn. Sci. https://doi.org/10.1016/S1364-6613(00)01769-1

Lien, M.-C., Ruthruff, E., Johnston, J.C., 2006. Attentional Limitations in Doing Two Tasks at Once The Search for Exceptions.

Marois, R., Ivanoff, J., 2005. Capacity limits of information processing in the brain. Trends Cogn. Sci. 9, 296–305. https://doi.org/10.1016/j.tics.2005.04.010

Meiran, N., Gotler, A., 2001. Modelling cognitive control in task switching and ageing. Eur. J. Cogn. Psychol. 13, 165–186. https://doi.org/10.1080/09541440042000269

Meiran, N., Gotler, A., Perlman, A., 2001. Old age is associated with a pattern of relatively intact and relatively impaired task-set switching abilities. Journals Gerontol. -Ser. B Psychol. Sci. Soc. Sci. 56, P88–P102. https://doi.org/10.1093/geronb/56.2.P88

Melis, A., Soetens, E., Van Der Molen, M.W., 2002. Process-specific slowing with advancing age: Evidence derived from the analysis of sequential effects. Brain Cogn. https://doi.org/10.1006/brcg.2001.1508

Meyer, D.E., Kieras, D.E., 1997a. A Computational Theory of Executive Cognitive Processes and Multiple-Task Performance: Part 2. Accounts of Psychological Refractory-Period Phenomena. Psychol. Rev. 104, 749–791. https://doi.org/10.1037/0033-295X.104.4.749

Meyer, D.E., Kieras, D.E., 1997b. A Computational Theory of Executive Cognitive Processes and Multiple-Task Performance: Part 1. Basic Mechanisms. Psychol. Rev. 104, 3–65. https://doi.org/10.1037/0033-295X.104.1.3

Mirelman, A., Herman, T., Brozgol, M., Dorfman, M., Sprecher, E., Schweiger, A., Giladi, N., Hausdorff, J.M., 2012. Executive function and falls in older adults: New findings from a five-year prospective study link fall risk to cognition. PLoS One 7, 1–8. https://doi.org/10.1371/journal.pone.0040297

Nasreddine, Z.S., Phillips, N.A., Bédirian, V., Charbonneau, S., Whitehead, V., Collin, I., Cummings, J.L., Chertkow, H., 2005. The Montreal Cognitive Assessment, MoCA: A brief screening tool for mild cognitive impairment. J. Am. Geriatr. Soc. https://doi.org/10.1111/j.1532-5415.2005.53221.x

Navon, D., Miller, J., 2002. Queuing or sharing? A critical evaluation of the single-bottleneck notion. Cogn. Psychol. 44, 193–251. https://doi.org/10.1006/cogp.2001.0767

Park, D.C., Polk, T.A., Park, R., Minear, M., Savage, A., Smith, M.R., 2004. Aging reduces neural specialization in ventral visual cortex. Proc. Natl. Acad. Sci. U. S. A. 101, 13091–13095. https://doi.org/10.1073/pnas.0405148101

Park, Seo, J., Kim, Y., Ko, M., 2014. Long-term effects of transcranial direct current stimulation combined with computer-assisted cognitive training in healthy older adults. Neuroreport 25, 122–126. https://doi.org/10.1097/WNR.0000000000000080

Pashler, H., 1994. Dual-task interference in simple tasks: Data and theory. Psychol. Bull. 116, 220–244. https://doi.org/10.1037/0033-2909.116.2.220

Pashler, H., 1984. Processing stages in overlapping tasks: Evidence for a central bottleneck. J. Exp. Psychol. Hum. Percept. Perform. 10, 358–377. https://doi.org/10.1037/0096-1523.10.3.358

Payer, D., Marshuetz, C., Sutton, B., Hebrank, A., Welsh, R.C., Park, D.C., 2006. Decreased neural specialization in old adults on a working memory task. Neuroreport 17, 487–491. https://doi.org/10.1097/01.wnr.0000209005.40481.31

Persson, N., Ghisletta, P., Dahle, C.L., Bender, A.R., Yang, Y., Yuan, P., Daugherty, A.M., Raz, N., 2016. Regional brain shrinkage and change in cognitive performance over two years: The bidirectional influences of the brain and cognitive reserve factors. Neuroimage 126, 15–26. https://doi.org/10.1016/j.neuroimage.2015.11.028

Salthouse, T.A., Miles, J.D., 2002. Aging and time-sharing aspects of executive control. Mem. Cogn. 30, 572–582. https://doi.org/10.3758/BF03194958

Sanders, A.F., 2013. Elements of Human Performance, Elements of Human Performance. Psychology Press. https://doi.org/10.4324/9780203774250

Sheikh, J.I., Yesavage, J.A., 1986. Geriatric Depression Scale (GDS) Jerome. Clin. Gerontol. 5, 165–173. https://doi.org/10.1300/J018v05n01

Sigman, M., Dehaene, S., 2008. Brain mechanisms of serial and parallel processing during dual-task performance. J. Neurosci. 28, 7585–7598. https://doi.org/10.1523/JNEUROSCI.0948-08.2008

Smith, A., 2011. Symbol Digit Modalities Test, in: Encyclopedia of Clinical Neuropsychology. Springer New York, New York, NY, pp. 2444–2444. https://doi.org/10.1007/978-0-387-79948-3_6121

Stelmach, G.E., Goggin, N.L., García-Colera, A., 1987. Movement specification time with age. Exp. Aging Res. 13, 39–46. https://doi.org/10.1080/03610738708259298

Stephens, J.A., Berryhill, M.E., 2016. Older Adults Improve on Everyday Tasks after Working Memory Training and Neurostimulation. Brain Stimul. 9, 553–559. https://doi.org/10.1016/j.brs.2016.04.001

Tombu, M., Asplund, C., Dux, P.E., Godwin, D., Martin, J.W., Marois, R., 2011. A unified attentional bottleneck in the human brain. Proc. Natl. Acad. Sci. U. S. A. 108, 13426–13431. https://doi.org/10.1073/pnas.1103583108

Tombu, M., Jolicoæur, P., 2003. A Central Capacity Sharing Model of Dual-Task Performance. J. Exp. Psychol. Hum. Percept. Perform. 29, 3–18. https://doi.org/10.1037/0096-1523.29.1.3

Unsworth, N., Heitz, R.P., Schrock, J.C., Engle, R.W., 2005. An automated version of the operation span task. Behav. Res. Methods 37, 498–505. https://doi.org/10.3758/BF03192720

van Doorn, J., van den Bergh, D., Bohm, U., Dablander, F., Derks, K., Draws, T., Evans, N.J., Gronau, Q.F., Hinne, M., Kucharský, Š., Ly, A., Marsman, M., Matzke, D., Raj, A., Sarafoglou, A., Stefan, A., Voelkel, J.G., Wagenmakers, E.-J., 2019. The JASP Guidelines for Conducting and Reporting a Bayesian Analysis. PsyArxiv Prepr. 0–31. https://doi.org/10.31234/osf.io/yqxfr

Verhaeghen, P., 2011. Aging and executive control: Reports of a demise greatly exaggerated. Curr. Dir. Psychol. Sci. 20, 174–180. https://doi.org/10.1177/0963721411408772

Verhaeghen, P., Cerella, J., 2002. Aging, executive control, and attention: a review of meta-analyses. Neurosci. Biobehav. Rev. 26, 849–857. https://doi.org/10.1016/S0149-7634(02)00071-4

Verhaeghen, P., Steitz, D.W., Sliwinski, M.J., Cerella, J., 2003. Aging and dual-task performance: A meta-analysis. Psychol. Aging. https://doi.org/10.1037/0882-7974.18.3.443

Welford, A.T., 1952. the ‘Psychological Refractory Period’ and the Timing of High‐Speed Performance—a Review and a Theory. Br. J. Psychol. Gen. Sect. 43, 2–19. https://doi.org/10.1111/j.2044-8295.1952.tb00322.x

